# Modulation of exploration-avoidance behaviors by vmPFC-projecting BLA neurons

**DOI:** 10.64898/2026.03.01.708792

**Authors:** Huixin Huang, Shreesh P. Mysore, Hita Adwanikar

## Abstract

The ventromedial prefrontal cortex (vmPFC) is known to regulate exploration-avoidance behaviors. Separately, the basolateral amygdala (BLA), which plays a pivotal role in interpreting the emotional valence of external stimuli, is known to send information about aversive cues to the mPFC. Here, we investigated the role of vmPFC-projecting BLA neurons in modulating exploration-avoidance behaviors. We selectively and reversibly inhibited this subset of BLA neurons using a projection-defined chemogenetic approach in mice engaged in a classic unconditioned exploration-avoidance task, the elevated zero maze (EZM). Saline injection served as a control for administration of the chemogenetic ligand CNO. We found that inhibiting vmPFC-projecting BLA neurons with CNO injections promotes avoidance behaviors. This effect could not be attributed to the effects of injection. It could also not be attributed to repeated maze exposure because, whereas we observed behavioral habituation in naive mice following repeated exposure at 1 hr, in mice mimicking treatment manipulation (IP injection), no habituation was observed at this timepoint. Together, our results reveal that vmPFC-projecting BLA neurons promote safety rather than aversive signals in the context of unconditioned exploration-avoidance behavior. Surprisingly, these findings are directly opposed to previous work that suppressed BLA projection fibers in the mPFC. These contrasting results suggest pathway-dependent roles of mPFC-projecting BLA neurons (direct axonal vs. complex somatic, due to direct + indirect), or alternatively, specializations in the roles of mPFC subregions (vmPFC vs. dmPFC) in exploration-avoidance behavior. Our results also provide guidance for future experimental designs involving repeated exposure to the EZM.

## Introduction

Rodents exhibit an innate conflict between the drive to explore novel environments and the need to avoid open or brightly lit spaces that may signal predation risk. This exploration-avoidance conflict is widely regarded as a core behavioral correlate of anxiety in animals (La-Vu et al., 2020). To quantify this behavior in mice, researchers commonly use paradigms such as the elevated plus maze, elevated zero maze, open field test, and light-dark box (Calhoon & Tye, 2015; La-Vu et al., 2020; Tovote et al., 2015). These tasks leverage rodents’ natural preference for enclosed, dark spaces over open, illuminated areas, thereby inducing a measurable conflict between exploration and avoidance tendencies.

The basolateral amygdala (BLA) and the medial prefrontal cortex (mPFC), two anatomically and functionally interconnected regions, are central to modulating exploration-avoidance behavior. The BLA plays a pivotal role in interpreting the emotional valence of external stimuli, forwarding this information via diverse pathways, with the medial prefrontal cortex (mPFC) being a prominent target (Zhang & Li, 2018). The mPFC receives and evaluates cues from the BLA (as well as behaviorally relevant contextual information from the ventral hippocampus), and provides feedback to downstream regions, including the BLA itself, the striatum, and the nucleus accumbens, to either enable or suppress avoidant responses (Calhoon & Tye, 2015). As such, the BLA-mPFC circuit constitutes a key node for coding aversive cues and regulating exploration-avoidance behaviors.

The BLA projects to the mPFC via both direct and indirect pathways, including through the ventral hippocampus (vHPC) (Calhoon & Tye, 2015; Tovote et al., 2015). Direct BLA projections to the mPFC are implicated in responses to aversive cues during fear conditioning (Herry et al., 2008; Laviolette et al., 2005) and have been shown to modulate anxiety-related behaviors (Felix-Ortiz et al., 2016).Previous work has primarily focused on BLA projections to the dorsal mPFC (dmPFC) and identified an anxiogenic effect of this pathway (Felix-Ortiz et al., 2016). However, recent studies suggest functional heterogeneity within the mPFC, the dmPFC and the ventral mPFC (vmPFC), in the context of fear conditioning: the dmPFC supports fear expression, whereas the vmPFC promotes fear extinction (Burgos-Robles et al., 2009; Capuzzo & Floresco, 2020; Do-Monte et al., 2015; Klavir et al., 2017; Sierra-Mercado et al., 2011). These subregional differences raise the possibility that BLA projections to the vmPFC may play a distinct, potentially anxiolytic role in exploration-avoidance conflict. However, this pathway has not been directly interrogated.

To address these gaps, we used projection-defined chemogenetic approach to selectively inhibit BLA neurons that project to the vmPFC in freely behaving mice engaged in an exploration-avoidance task (elevated zero maze; EZM). Because this experiment requires repeated testing across hours and days, we first assessed whether repeated exposure to EZM at various inter-trial intervals, or treatment intervention (in this case, IP injection), induced behavioral changes that could confound our interpretation. We ran a separate cohort of animals on the EZM and compared their behaviors at multiple time points corresponding to the experimental conditions with CNO/saline. Based on these results, we optimized our experimental design to minimize potential carryover effects. Beyond guiding experimental design for studies examining the effects of drug injections (e.g., chemogenetic manipulation), our findings on the impact of repeated maze exposure may also inform broader studies aimed at investigating the neural mechanisms underlying stability of behavioral and neural measures across time (Johnson et al., 2022; Huang et al., 2026).

## RESULTS

### Repeated EZM exposure induces decreased exploration after 1 hour, but not over longer timescales

To determine the optimal timescale for our BLA-vmPFC chemogenetic manipulation, we first assessed how repeated exposure to the EZM affects exploration-avoidance behavior across intervals ranging from hours to weeks (1 hour, 1 day, 1 week, and 2 weeks), in a cohort of 10 mice.

The experimental procedure was as follows (Fig. 1A). Mouse was first acclimated to the behavioral room for 30 minutes in its home cage. Then the mouse would freely explore the EZM for 20 minutes. For the 1-hour condition, the mouse was returned to the EZM for a second 20-minute session one hour after the first. For the 1-day, 1-week, and 2-week conditions, animals underwent the same acclimation and 20-minute EZM session after the specified delay. To quantify exploration-avoidance behavior, we measured the percentage of time the mouse’s head (rather than its body centroid; see ref: Huang et al., 2026) occupied the open arms of the maze.

**Figure 1.**
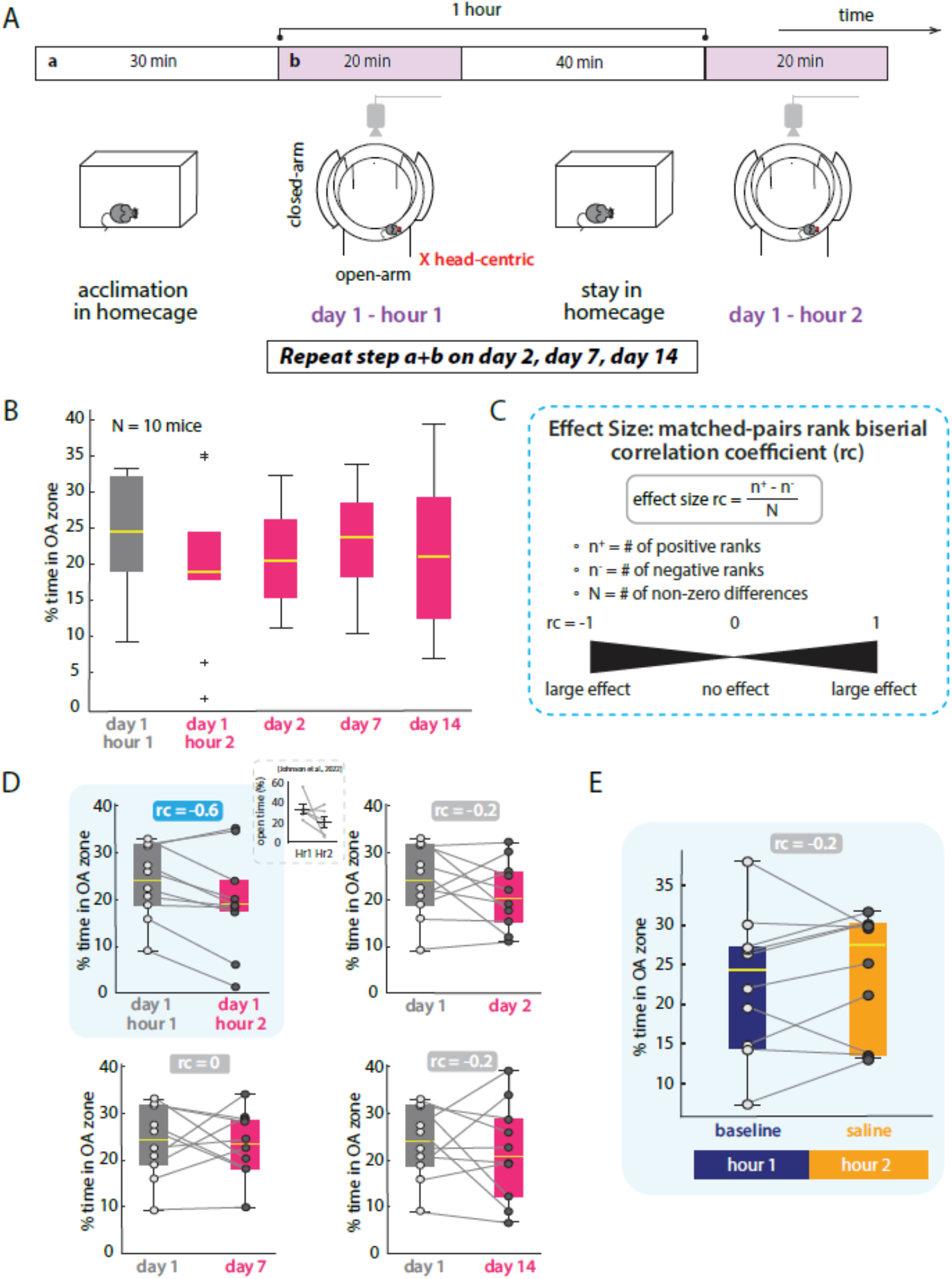
Behavioral effects of repeated EZM exposure. **(A)** Schematic of experimental design. Mice (n = 10) were exposed to EZM multiple times, with inter-session intervals ranging from 1 hour to 14 days. Each session was preceded by a 30-minute habituation period in the behavioral room. **(B)** Time spent in the open arm (OA) of the EZM across five time points: initial exposure (day 1 hour 1), repeated exposure after 1 hour (day 1 hour 2), and subsequent sessions on days 2, 7, and 14. **(C)** Illustration of the matched-pairs rank-biserial correlation coefficient (rc), used to quantify direction and magnitude of behavioral change across conditions. Positive rc indicates increased exploration; negative rc indicates reduced exploration. **(D)** Pairwise comparisons between the initial EZM session (day 1 hour 1) and each subsequent session. The rc is shown for each pair. Top-left inset is adapted from Johnson et al. (2022). **(E)** Time spent in the OA during a baseline EZM session, and a repeated session conducted 1 hour later following intraperitoneal saline injection.

We conducted a nonparametric Friedman’s test on behavior across all time points: day 1 hour 1, day 1 hour 2, day 2, day 7, and day 14. This analysis did not reveal a significant main effect of the time condition (p=0.56; Fig. 1B). To further examine potential short-term changes, we performed pairwise comparisons between the first exposure (day 1 hour 1) and each subsequent time point. After a 1-hour delay, 8 out of 10 animals exhibited reduced exploratory behavior, as reflected by decreased time spent in the open arms. To quantify these changes, we calculated the matched-pairs rank biserial correlation coefficient (rc) (Cureton, 1956), which captures effect size by comparing the number of positive (increase in) versus negative (decrease in) behavioral changes between paired conditions. Values of rc range from −1 (consistent decrease) to +1 (consistent increase), with 0 indicating no net change (Fig. 1C). The rc between hour 1 and hour 2 was −0.6, indicating a moderate decline in exploratory tendency (Fig. 1D, top-left).

In contrast, comparison between hour 1 and later time points (1 day, 1 week, and 2 weeks) yielded rc values near zero, indicating stable behavior with no consistent trend toward increased or decreased exploration (Fig. 1D, top-right, bottom). We conclude that repeated exposure to the EZM leads to a transient suppression of exploration after one hour, but not after one day.

### Repeated EZM exposure combined with injection has no effect after 1 hour

Our chemogenetic experiments involve intraperitoneal (IP) injection of clozapine-N-oxide (CNO), which introduces an additional potential confounding factor: the interaction between injection and repeated maze exposure could impact animals’ behavior in unexpected way. To test this, we replicated the 1-hour interval experiment described above, with one modification - mice received an IP injection of saline immediately after the first EZM session.

We compared open-arm exploration between the initial session (hour 1/baseline) and the post-injection session (hour 2/saline). The matched-pairs rank biserial correlation coefficient (rc) was 0.2, indicating a minimal change in behavior (Fig. 1E).

This result suggests that, although repeated exposure alone tends to suppress exploratory behavior after 1 hour, the addition of injection appears to mitigate this effect. Thus, the combined effect of repeated exposure plus injection produces little net behavioral change on a 1-hour scale. We conclude that for experiments involving both EZM re-testing and injection (e.g., CNO administration), a 1-hour interval between sessions is acceptable and unlikely to confound interpretation.

### Chemogenetic inhibition of vmPFC-projecting BLA neurons reduces exploration

To determine the role of BLA neurons projecting to the vmPFC in exploration-avoidance behavior, we used a projection-specific chemogenetic approach. A retrograde Cre-expressing virus (Ef1a-mCherry-IRES-Cre) was injected into the right ventromedial mPFC (vmPFC), and 2–4 weeks later, a Cre-dependent inhibitory DREADD virus (AAV-DIO-hM4Di) was injected into the ipsilateral BLA. Following an additional 2–4 weeks to allow for sufficient viral expression, mice were tested on the elevated zero maze (EZM) to assess the effect of chemogenetic inhibition (Fig. 2A, top).

**Figure 2.**
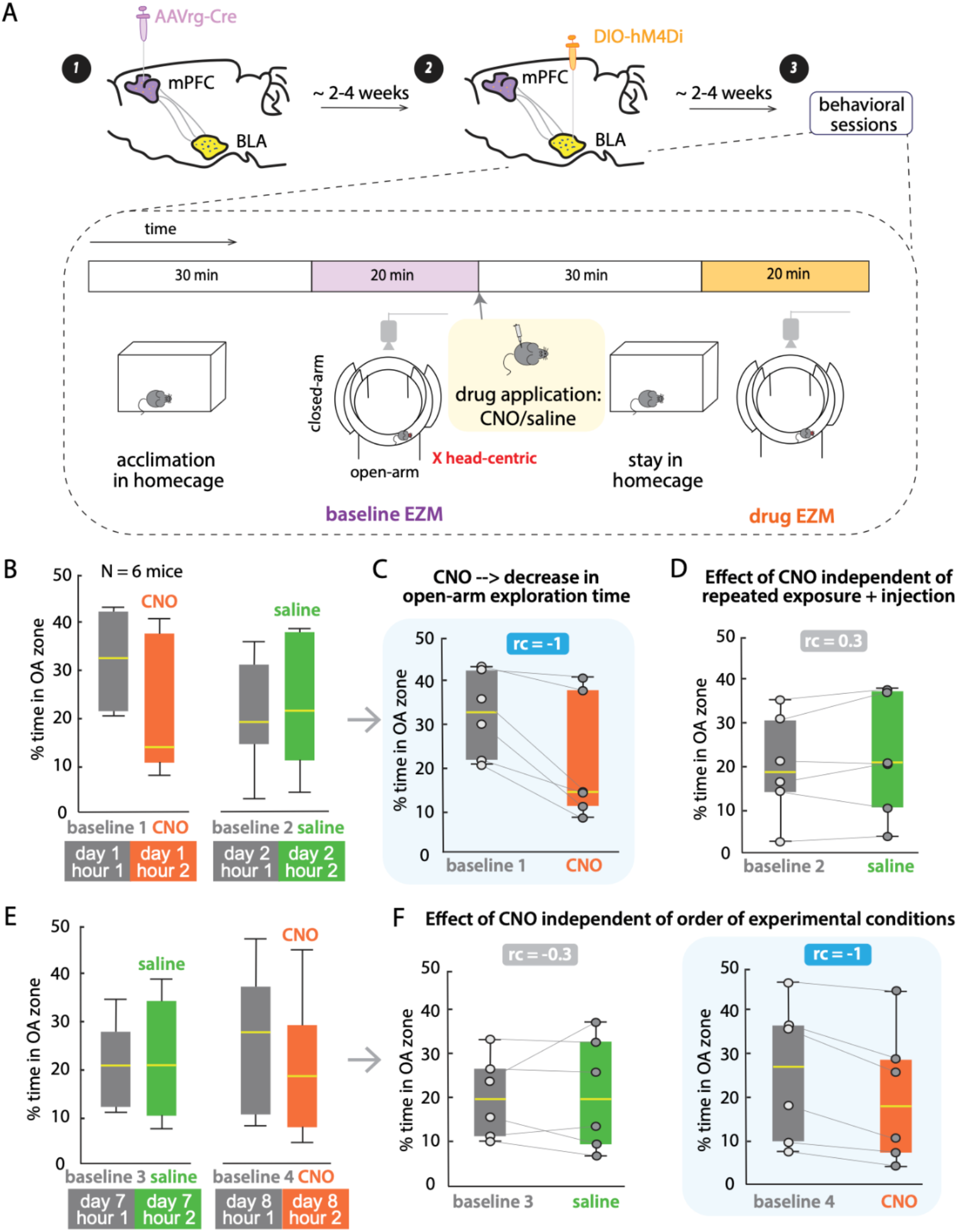
Chemogenetic inhibition of vmPFC-projecting BLA neurons reduces exploration in the EZM. **(A)** Experimental design for projection-defined chemogenetic inhibition. (Top) A retrograde Cre-expressing virus (Ef1a-mCherry-IRES-Cre) was infused into the vmPFC, followed 2-4 weeks later by a Cre-dependent DREADD virus (AAV-DIO-hM4Di) into the ipsilateral BLA (n = 6). (Bottom) Schematic of behavioral sessions. Each mouse underwent a 30-minute habituation period, followed by a 20-minute baseline EZM session. Immediately afterward, mice received an intraperitoneal injection of CNO or saline. 30 minutes post-injection, they were re-exposed to the EZM for an additional 20-minute session. **(B)** Summary of percentage time spent in the OA across four sessions: baseline 1 (day 1 hour 1), CNO session (day 1 hour 2), baseline 2 (day 2 hour 1), and saline session (day 2 hour 2). **(C)** Derived from (B). Pairwise comparisons of OA exploration between baseline 1 and the CNO session. **(D)** Derived from (B). Pairwise comparisons between baseline 2 and the saline session, controlling for the combined effects of repeated exposure and injection. **(E)** Summary of OA exploration in a counterbalanced order: baseline 3 (day 7 hour 1), saline session (day 7 hour 2), baseline 4 (day 8 hour 1), and CNO session (day 8 hour 2). **(F)** Derived from (E). Pairwise comparisons between baseline and saline/CNO sessions in the counterbalanced order, controlling for the effect of order of experimental conditions.

Each mouse was acclimated to the behavioral room for 30 minutes in its home cage, followed by a 20-minute baseline EZM session. Based on our earlier findings that a combination of injection and repeated exposure does not alter exploratory behavior at the 1-hour interval, we administered CNO intraperitoneally immediately after the baseline session to inhibit mPFC-projecting BLA neurons. Mice were then returned to the EZM for a second 20-minute session approximately 1 hour after the start of the baseline session (Fig. 2A, bottom).

We observed a consistent reduction in open-arm exploration across all animals following CNO administration, reflected by a rank biserial correlation coefficient (rc) of −1 (Fig. 2B, left; Fig. 2C). This indicates a robust suppression of exploratory behavior when vmPFC-projecting BLA neurons are inhibited. These findings suggest that this projection supports approach behavior and may reduce avoidant behavior in open, potentially threatening environments.

### The effect of CNO is independent of repeated exposure + injection and the order of experimental conditions

To confirm that the observed behavioral effect was not due to repeated exposure to the EZM combined with injection alone, we performed a within-subject control experiment in the same cohort. On the day following the baseline-CNO session, mice underwent a baseline-saline session in which the procedure was identical to the baseline-CNO session, except that CNO was replaced with saline (Fig. 2A). The matched-pairs rank biserial correlation coefficient (rc) for the baseline-saline comparison was 0.3, indicating minimal behavioral change and supporting the conclusion that injection and repeated exposure alone do not account for the reduction in exploration (Fig. 2D).

Because the CNO session always preceded the saline session in the initial experiment, we next asked whether the order of conditions might have contributed to the effect. One week later, we repeated both sessions in reverse order: mice first underwent the baseline-saline session, followed by the baseline-CNO session on the next day. The results replicated our original findings - saline had no consistent effect, while CNO administration led to a robust decrease in exploration across all animals (Fig. 2E-F).

Together, these control experiments demonstrate that the suppression of exploratory behavior is specifically driven by chemogenetic inhibition of mPFC-projecting BLA neurons. The behavioral effect is not attributable to repeated maze exposure, IP injection, or the order in which experimental conditions were presented.

## Discussion

In summary, our findings demonstrate that chemogenetic inhibition of BLA neurons projecting to the vmPFC promotes avoidance behavior in mice, as evidenced by reduced exploration in the EZM. Crucially, this effect cannot be attributed to confounding factors such as repeated exposure to the maze, IP injection, or the order of experimental conditions, as verified by multiple within-subject control experiments. Additionally, we observed that while repeated exposure to EZM induces behavioral adaptation within an hour (unlike previous work: Johnson et al., 2022), this effect dissipates by 24 hours. Interestingly, when repeated exposure was combined with injection after one hour, no behavioral change was observed, as shown by two independent experiments.

Our task design was carefully chosen to ensure validity over the comparisons required. Since behavioral habituation was minimal in the 1-hour repeated exposure following an injection, behavior in CNO, or saline control, conditions was measured on the same day as baseline, with an inter-session-interval of 1 hour. To avoid additional confounds due to multiple IP injections and EZM sessions an hour apart, we did not run saline-control and CNO conditions on the same day. Separately, counterbalancing the order of CNO treatment with saline control allowed us to confirm that order of treatment does not influence the finding of reduced exploratory behavior when vmPFC projecting BLA neurons are inhibited.

Surprisingly, our results are directly opposed to prior work using optogenetics to suppress BLA projection fibers in the mPFC, which reported anxiolytic effects and increased open arm exploration (Felix-Ortiz et al., 2016). In our study, inhibiting combined direct and indirect BLA-mPFC projection by suppressing vmPFC-projecting BLA neurons completely reversed the previous finding and reduced exploration of the open arm. A key difference between our study and this previous work could be the specificity of the sub-region target. While the coordinates used in Felix-Ortiz et al. (2016) targeted inactivation of BLA terminals in more dorsal mPFC neurons, our method targeted BLA input to a more ventral mPFC population. Growing evidence suggests functional differences between dmPFC and vmPFC, the two subregions of mPFC, in the context of fear conditioning: dmPFC supports fear expression whereas vmPFC contributes to fear extinction and safety learning (Burgos-Robles et al., 2009; Capuzzo & Floresco, 2020; Do-Monte et al., 2015; Klavir et al., 2017; Sierra-Mercado et al., 2011). This suggests that dmPFC enables aversive interpretations of environmental cues while vmPFC generates safety signals that attenuate aversive responses in fear learning and anxiety regulation. In this context, it has been suggested that BLA projections to mPFC are also important in coding threatful cues and in modulating expression of aversive states (Herry et al., 2008; Laviolette et al., 2005). Remarkably, amygdala neurons that fire to threatening cues have been found to preferentially project to the dmPFC, while those active after extinction of conditioned fear preferentially target the vmPFC (Senn et al., 2014). Together, these results suggest not only that functional activation of amygdala afferents can activate the corresponding target mPFC subdivision (Likhtik & Paz, 2015), but also that it will evoke differential effects on anxiety-like behavior.

Another possible explanation could be that the contrasting results between cell-body versus axonal manipulations of this this subgroup of BLA neurons suggests complex, pathway-dependent role in exploration-avoidance behavior. It is possible that direct and indirect BLA-mPFC projections serve distinct yet complementary functions in modulating exploration; inhibiting direct BLA-mPFC pathway (by targeting the axonal terminals) could remove the rapid communication of emotional valence and threat information, and produce anxiolytic-like behavioral effects. On the other hand, BLA neurons might also indirectly project to mPFC through regions like vHPC, in which the valence information is integrated with contextual or spatial information encoded by vHPC before reaching mPFC. Thus reduced activity in this population of BLA neurons could contribute to a more flexible, state-dependent modulation of exploration-avoidance behavior. However, the degree to which the direct and indirect BLA-mPFC pathway work in a synergistic or complementary manner is unclear.

This anatomical and functional specificity likely contributes to the divergent behavioral outcomes observed across studies. Our finding that BLA-vmPFC inhibition promotes avoidance behavior is consistent with the hypothesis that this pathway facilitates safety signaling and approach behavior in anxiogenic environments.

Our results also have important methodological implications for behavioral assays involving repeated EZM testing. Although EZM is widely used to assess exploration-avoidance behavior, repeated exposure can induce habituation/behavioral adaptation, potentially confounding studies examining the effects of drug injections (e.g., chemogenetic manipulation) on exploration-avoidance behaviors or the neural mechanisms underlying behavioral stability. Previous studies have reported the effects of repeated exposure to the EZM (Popovitz et al., 2019; Johnson et al., 2022), but there is a lack of clarity for the short timescale (hours and days). This is particularly relevant for experiments involving pharmacological intervention, which often require repeated behavioral testing within short timescales. Our findings clarify that a reduction in exploratory behavior can occur within an hour of exposure, but returns to baseline across days. Moreover, we demonstrate that combining injection with repeated exposure ameliorates this change, supporting a 1-hour inter-session interval for drug injection (e.g. chemogenetic manipulation) without introducing behavioral artifacts. These results are particularly useful as a growing number of studies become interested in investigating the neural representation underlying exploration-avoidance behavior (Huang et al., 2026). Recent work suggests that while single neural code may be unstable, population-level neural codes stably represent exploration-avoidance states across time (Johnson et al., 2022). Behavioral stability is therefore critical for probing these underlying codes. Our results indicate that both repeated exposure and other treatment manipulations can affect how the brain supports behavioral stability in EZM.

Beyond the behavior effects of manipulating vmPFC-projecting BLA neurons, it will be interesting to also study how vmPFC neural dynamics are modulated by BLA input to drive behavioral change. Simultaneous electrophysiological recording/calcium imaging in vmPFC paired with chemogenetic/optogenetic perturbation of BLA inputs will be useful for dissecting the underlying mechanisms. It is possible that disrupting vmPFC-projecting BLA neurons promotes avoidance by different potential mechanisms; such as by increasing firing rates of individual vmPFC neurons specifically in aversive zones, by increasing the proportion of vmPFC neurons that preferentially encode aversive signals, or by altering the structure of ensemble trajectories in population state space. Such experiments would provide mechanistic insight into how BLA-mPFC communication regulates the balance between exploration and avoidance in potentially threatening environments.

## MATERIALS AND METHODS

### Subjects

A total of sixteen mice were used in this study. The repeated exposure tests were conducted on five male and five female C57BL/6 mice aged 8-10 weeks and BLA-mPFC experiments were conducted on three male and three female C57BL/6 mice aged 3-6 months obtained from Jackson Laboratories, Bar Harbor, Maine. Mice for repeated exposure tests were group-housed and for BLA-mPFC experiments were single-housed in behaviorally enriched home cages. The mice were allowed to acclimatize to the animal care facility for at least a week, under a 12-hour light cycle (7am to 7pm) with constant temperature (22°C) and humidity (40%), before starting any testing procedures. Food and water were provided ad libitum throughout the experiments. Experiments occurred during the standard light cycle. All protocols and animal care were in accordance with NIH guidelines for care and use of laboratory animals and approved by the Johns Hopkins University Institutional Animal Care and Use Committee.

### Surgery

We performed surgery and virus injections on six mice aged 3–6 months. Briefly, each mouse was allowed to acclimate to the surgery room for 30 minutes before being placed in an anesthetic chamber (3% isoflurane, 1.5 L/min O_2_) for 5 minutes. The mouse was then secured in a stereotaxic frame (Kopf Instruments), and body temperature was maintained at 36.8 °C using a heating pad. Prior to the surgical procedure, the animal received an intraperitoneal injection of meloxicam (0.2 mL). A craniotomy was performed at the stereotaxic coordinates targeting the vmPFC. A 350 nL injection of retrograde Cre-expressing virus (Ef1a-mCherry-IRES-Cre) was delivered using a 0.5 mL microsyringe and motorized pump (Harvard Apparatus); the syringe needle was lowered at a rate of 200 µm/min into the right vmPFC (y = +1.86 mm AP, x = +0.25 mm ML, z = −2.80 mm DV relative to Bregma). Following the surgery, animals were allowed to recover under a heat lamp before being returned to the animal facility. Meloxicam (0.1 mL) was administered once daily for 3 days post-surgery. Mice were allowed to recover for 2-4 weeks to allow retrograde labeling of BLA neurons.

Subsequently, we performed a second surgery. All preparation and post-surgical care procedures were the same as described above. A second craniotomy was made at the coordinates targeting the BLA. A 250 nL injection of Cre-dependent DREADD virus (AAV-DIO-hM4Di) was delivered using the same syringe and motorized pump; the needle was lowered at 200 µm/min into the right BLA (y = −1.65 mm AP, x = +2.80 mm ML, z = −4.80 mm DV relative to Bregma). Mice recovered for an additional 2-4 weeks before undergoing behavioral testing.

### Behavioral essay: elevated zero maze

The elevated zero maze (EZM) consisted of a circular platform raised above ground level (6.1 cm width, 40 cm inner diameter, 72.4 cm above ground; Med Associates, St. Albans, VT, USA) and divided into four quadrants. Two opposite quadrants were enclosed by 20.3 cm high walls (“closed arms”), while the other two had no walls (“open arms”). A camera was centered above the maze apparatus to record mouse behavior. The apparatus was placed within an area surrounded by thick black curtains, and lights were dimmed. Following a 30-minute acclimation to the experimental room, animals were placed in a designated open arm of the EZM, and their freely moving behavior was recorded for 20 minutes. An automated behavioral monitoring system (EthoVision, Noldus Inc., v11.5; frame rate 8.31 Hz) was used to track the animal’s head-centroid motion trajectories from the recorded videos. The animal was returned to its home cage and the animal facility at the end of the experiment.

### Experimental procedure

Described in detail in and around figure 1A and figure 2A.

### Statistics

All statistical tests were performed in MATLAB. Friedman’s test (the nonparametric equivalent of repeated-measure ANOVA) was used with time condition as the independent variable. In all statistical tests, the significance threshold was set at p < 0.05. To quantify behavioral changes, we used the matched-pairs rank biserial correlation coefficient (rc), a measure of effect size defined as the difference between the number of positive occurrences (increase in values from condition 1 to 2) and the number of negative occurrences (decrease in values), normalized by the total number of paired comparisons. Values of rc range from −1 (consistent decrease) to +1 (consistent increase), with 0 indicating no net change.

## ACKNOWLEDGMENTS

We thank members of the Adwanikar and Mysore groups for feedback and discussions.

## AUTHOR CONTRIBUTIONS

HA and SPM conceived the project. HH performed the experiments, the analyses, and generated the figures. HH, HA and SPM wrote the paper.

